# Patterns of community science data use in peer-reviewed research on biodiversity

**DOI:** 10.1101/2022.10.17.512636

**Authors:** A.D. Binley, J.G. Vincent, T. Rytwinski, C.A. Proctor, E.S. Urness, S.A. Davis, P. Soroye, J.R. Bennett

**Affiliations:** Department of Biology, Carleton University, 1125 Colonel By Drive, Ottawa, ON, K1S 5B6, Canada; Canadian Centre for Evidence-Based Conservation, Institute of Environmental and Interdisciplinary Science, Carleton University, Ottawa, ON, K1S 5B6, Canada; Department of Biology, University of Ottawa, 75 Laurier Avenue E., Ottawa, ON, K1N 6N5, Canada

**Keywords:** big data, biodiversity, citizen science, conservation, evidence synthesis

## Abstract

Community science (“citizen science”) represent a potentially abundant and inexpensive source of information for biodiversity research. However, analyzing such data has inherent challenges. To explore where and how community science data are translated into scientific knowledge, we conducted a literature review in a sample of 334 peer-reviewed scientific articles. Specifically, we investigated how the use of community science data varied among taxonomic groups and geographic regions, and what threats to biodiversity, if any, were examined. Community science data were used mostly for research on birds and invertebrates, and the data used were mainly from the United States and the United Kingdom. Literature in certain countries used a wider breadth of projects, while others made repeated use of comparably fewer datasets. Community science efforts were largely used to measure abundance, trends, distributions, and range shifts. However, few articles linked these metrics to any particular threats to biodiversity. Furthermore, community science data were used infrequently for research on threatened species and limited mostly to count data rather than collecting more specific information such as life history, phenological or genetic data, suggesting that community science may be underutilized for these key aspects of biodiversity conservation. We conclude that even with the rise of community science data use in research, there remains tremendous potential to better use these existing datasets for biodiversity research.

## 1. Introduction

Understanding and responding to the biodiversity crisis requires extensive analysis of biodiversity data. Monitoring the world’s biodiversity however is no easy task, and much is still unknown about the patterns of species in space and time. In addition, monitoring efforts are biased both geographically (Nerlekar et al., 2019; Reddy & Dávalos, 2003; Yang et al., 2014) and taxonomically (Donaldson et al., 2017; Gordon et al., 2019), which in turn results in biased conservation efforts (Gordon et al., 2019).

Community science (or “citizen science”) is increasingly used to study biodiversity because it can often accomplish what conventional research cannot, namely the collection of long-term data across large spatial extents (Chandler et al., 2017). It can also be more cost effective at larger scales (Heigl et al., 2017), making it an appealing option for conservation initiatives with limited funding. In this paper, we define the term “community science” as the general public’s unpaid participation in biodiversity monitoring and data collection. Community science can take a variety of forms, from undirected programs recording opportunistic observations made by amateur naturalists (e.g. iNaturalist: Callaghan et al., 2020; eBird: Sullivan et al. 2014) to highly skilled volunteers operating under strict protocols and coordinated by government agencies (e.g. the Breeding Bird Survey (BBS); Hudson et al., 2017).

Although community science has proven to be a valuable resource, researchers may be reluctant to turn to these data for conservation biology research due to real and perceived limitations. The spatial and temporal scale of a community science project, along with scientific rigour and data availability, are major factors in predicting whether the data from a community science project are used in peer-reviewed literature (Theobald et al. 2015). Most data are concentrated around populous cities and accessible areas such as parks and near roads (Geldmann et al., 2016; Soroye et al., 2022) and are biased towards more charismatic species such as butterflies and birds (Theobald et al., 2015). Community science is often regarded only as a source of baseline surveillance data (Dickinson et al., 2010), and enlisting the public to help monitor sensitive species can pose significant risks (Soroye et al., 2022). These patterns could be perpetuated into the literature, compounding similar existing research biases (Donaldson et al., 2017).

Our primary goal with this evidence synthesis was to examine the patterns in community science use to illustrate potential biodiversity research opportunities. Broadly, we explored how community science data are being used in the peer-reviewed knowledge base, and how they could be better leveraged to inform conservation. To accomplish this, we searched and collated the peer-reviewed evidence base for articles that used community science to monitor terrestrial biodiversity, to ask the following specific questions: (1) how does community science use vary among taxonomic groups; (2) how does community science use vary among regions, and how does this variation interact with taxonomic patterns and country wealth; and (3) what threats to biodiversity are being addressed by research using community science? Though the availability of community science data has already been examined elsewhere (Chandler et al., 2017; Theobald et al., 2015), we build on this by investigating how community science data are used in the peer-reviewed biodiversity literature.

## 2. Methods

We were interested in peer-reviewed articles that used community science to collect data on terrestrial biodiversity. Our search methods were purposive, rather than systematic, as the goal was to obtain a representative sample of the prevalence of community science data in the published literature, rather than capture every article. We searched the academic databases ISI Web of Science Core Collections (WoSCC) and Scopus in January 2019. Our search strings were modified and refined iteratively through a scoping exercise which evaluated the sensitivity of search terms and associated wildcards. Search terms were kept intentionally general to avoid biasing our sample towards well-known community science projects (like the long-standing North American Breeding Bird Survey).

Our search string for WoSCC was as follows:

TS=((Crowdsourcing OR “community-based participatory research” OR “public participation” OR “participatory action research” OR “volunteer surveying” OR hunter OR naturalist OR atlas* OR Amateur OR hobbyist citizen OR “citizen researcher” OR “individual citizen scientist” OR collaborator OR “community researcher” OR contributor OR “lay knowledge holder” OR “general public” OR participant OR student OR pupil OR uncredentialled OR “non-credentialed researcher” OR nonacademic OR non-scientist OR volunteer) AND (ecolog* OR biodivers* OR conservation) AND (monitor* OR manage* OR survey) AND (species)) NOT (Marine OR restoration OR astron* OR medic*)

Our search string for Scopus was as follows:

TITLE-ABS-KEY ({community based participatory research} OR {community-based participatory research} OR “public participation” OR hunter OR naturalist OR atlas* OR amateur OR “citizen researcher” OR {individual citizen scientist} OR collaborator OR {lay knowledge holder}) AND TITLE-ABS-KEY (ecolog* OR biodivers* OR conservation) AND TITLE-ABS-KEY (monitor* OR manage* OR survey OR conserv*) AND TITLE-ABS-KEY (species) AND NOT TITLE-ABS-KEY (marine OR restoration OR astron* OR medic* OR cancer)

English search terms were used to conduct all searches in both databases. No language, date, or document type restrictions were applied during the search (i.e., if an abstract was in English but the article was in a different language), but only English language literature was included during the screening stage. Bibliographic databases were accessed using Carleton University’s institutional subscriptions (Appendix A).

Articles were screened at two stages (1) title and abstract, and (2) full-text, using pre-defined inclusion criteria. To be included in the analysis, articles were required to meet all criteria:

i. species were non-domesticated and had a terrestrial component to their lifecycle;
ii. members of the public (i.e., volunteers, both experts and non-experts) participated in the biodiversity monitoring and data collection; and
iii. the biological data collected by volunteers were used in analyses.

We did not include articles that solely outlined practices for successful citizen science programs or evaluated benefits to volunteers. In case of uncertainty, we screened the full text of the article for inclusion. Only articles written in English were retained, which we acknowledge is a limitation when examining regional patterns. Before independent screening began, a consistency check between four reviewers using a random subset of 50 articles was undertaken. The results of the consistency check were compared between reviewers, and all discrepancies were discussed to understand why an inclusion/exclusion decision was made. Revisions to the inclusion criteria were made as necessary.

We use the term ‘article’ to refer to an individual peer-reviewed document returned by our search, and ‘study’ to refer to a distinct species-location-time combination contained within an article. For data coding and extraction, a single article could contain multiple studies. For each study, we extracted information related to the community science project name(s), the species for which data were analyzed when included in the main body of the article, the geographic coverage at the country level, the type of threats examined, and the biological and ecological responses measured using community science data. For each article, we grouped the threats and responses into categories based on Dickinson et al. (2010) (Appendix A; Table S1). We categorized species as “threatened” if the authors stated that they were officially listed by a given conservation authority (such as a federal agency or the IUCN) as being at risk or in decline, or if the authors reported important population declines or other similar conservation concerns even if the species was not listed by a conservation authority.

To investigate how community science use varies among regions according to wealth, we used a linear model to examine the relationship between gross domestic product (GDP) per capita and community science data use in the literature. We used UN GDP per capita in USD, averaged between the years 1997 and 2019 inclusive for 194 countries (National Accounts Main Aggregates Database, 2021). We excluded overseas departments and territories from this analysis (e.g., French Guiana) as we could not always access GDP data for these locations. Taiwan and China were grouped together as the UN GDP data did not distinguish between the two countries. Data from Sudan and South Sudan were only available from 2008 – 2019.

The average GDP per capita was log transformed to better meet the assumptions of the linear model; however, there was still evidence of a relationship between the variance and the mean. Outliers were retained because the values were meaningful in this case (that is, GDP is inherently skewed with extreme values). The relationship between log-GDP per capita and the number of community science projects per country appeared to be linear.

To further explore geographical patterns of community science data use in the literature, we investigated the number of distinct community science projects in our sample by study location. Not all community science projects are well-suited or well-known enough to researchers for inclusion in a peer-reviewed publication. Some community science projects have been running for decades and vetted extensively, making them well-established sources of scientific data (e.g. the Breeding Bird Survey: Hudson et al. 2017), while newer and less structured programs can be of less certain quality (e.g., Kamp et al. 2016). Therefore, we were interested in gaining insight into whether more publications in each country correlated with a wider variety of data sources being used (within our sample), or whether the same established projects were being used repeatedly.

## 3. Results

### 3.1. Taxonomic patterns of community science data use in the literature

Across all 334 articles in our corpus, we found community science data were used to examine 739 species. Birds were the most prevalent taxon, with both the greatest number of articles (44.6%) that used community science data (Appendix A; Fig. S1), as well as the most individual species examined using these data (Fig. 1). Invertebrates were the second-most prevalent taxon, with 96 articles covering 70 unique families across 26 countries. Community science data on reptiles, amphibians and fungi were studied infrequently in our sample of the literature. Across taxa, species were less likely to be categorized as “threatened”, except for fungi.

**Figure 1.**
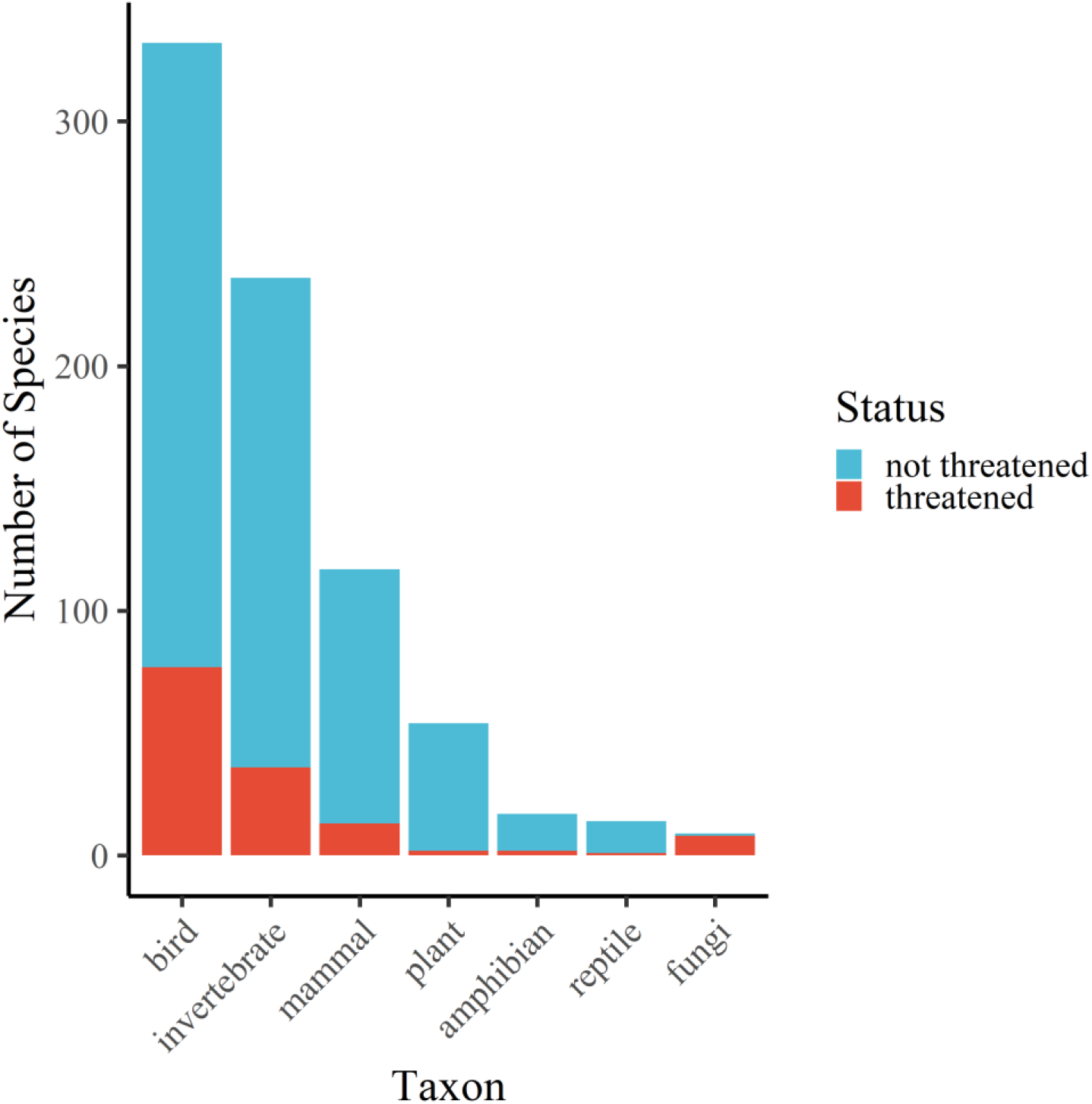
Number of unique species in each taxonomic group studied in the selection of articles captured by the literature review. Red represents the number of species that are described as “threatened” by the authors of the article, either due to population declines or other similar conservation concerns, or because they are officially listed under a given conservation authority (such as a federal agency or the IUCN) as being at risk or in decline. Species coded in blue were not reported to be in decline or to be listed by a conservation authority by the authors.

### 3.2. Geographic patterns of community science data use in the literature

In our sample, the articles making use of community science for biodiversity monitoring spanned 85 countries, with most located in the United Kingdom and the United States (Fig. 2). We found evidence of a log-linear relationship between country gross domestic product (GDP) per capita and the number of publications making use of community science data (β = 1.936, p < 0.0001, R^2^ = 0.108; Appendix B; Fig. S2). However, only a small proportion of the variation was explained by this relationship, and there are likely other factors at play that are unrelated to GDP.

**Figure 2.**
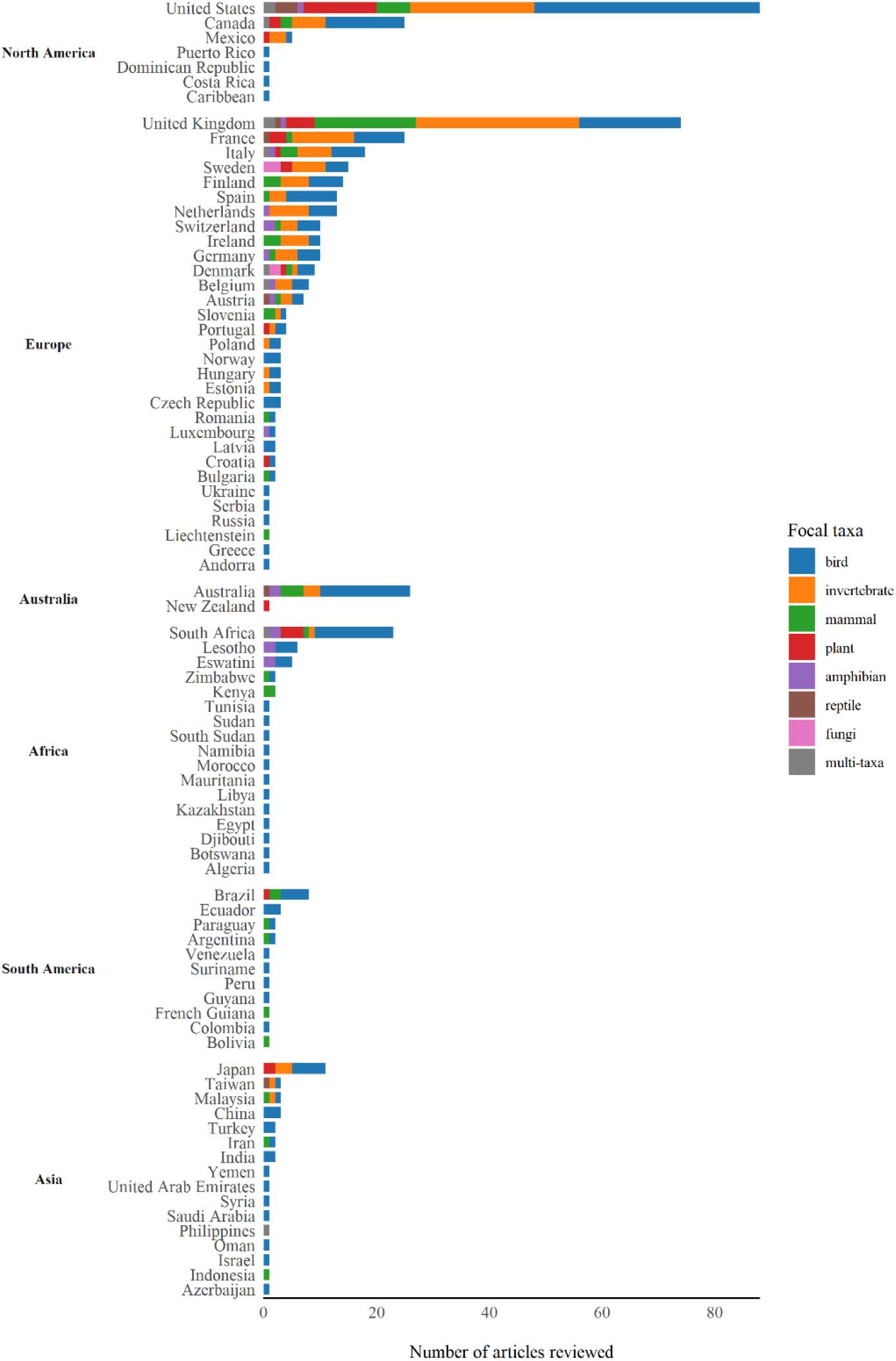
Number of articles in which biodiversity data were collected through community science, and the taxonomic breakdown of those articles, for each country where data were collected. “Multi-taxa” denotes an article or project that includes multiple taxonomic groups within a single article.

We found an almost 1:1 linear relationship between the number of reviewed articles conducted in each country and the total number of distinct community science projects that were used in publications in each country (β =0.91, R^2^ = 0.96, p = 3.36e-61, Fig. 3; see also Appendix B; Fig. S3 for patterns in residuals). That is, our sample suggests that the more publications using community science data in a country, the more likely these publications draw data from a breadth of available community science projects. We note that this is an underestimate of the number of actual community science projects in each country, given that we only included the projects found in our sample of peer-reviewed articles. Two of the more distinct outliers were Denmark and South Africa. Publications studying biodiversity in Denmark drew on a higher number of community science projects than expected based on the number of publications. In contrast, fewer distinct community science projects were used in South Africa than in Denmark, despite the former having more than double the number of community science publications (Fig. 3). We investigate these patterns further in Box 1.

**Figure 3.**
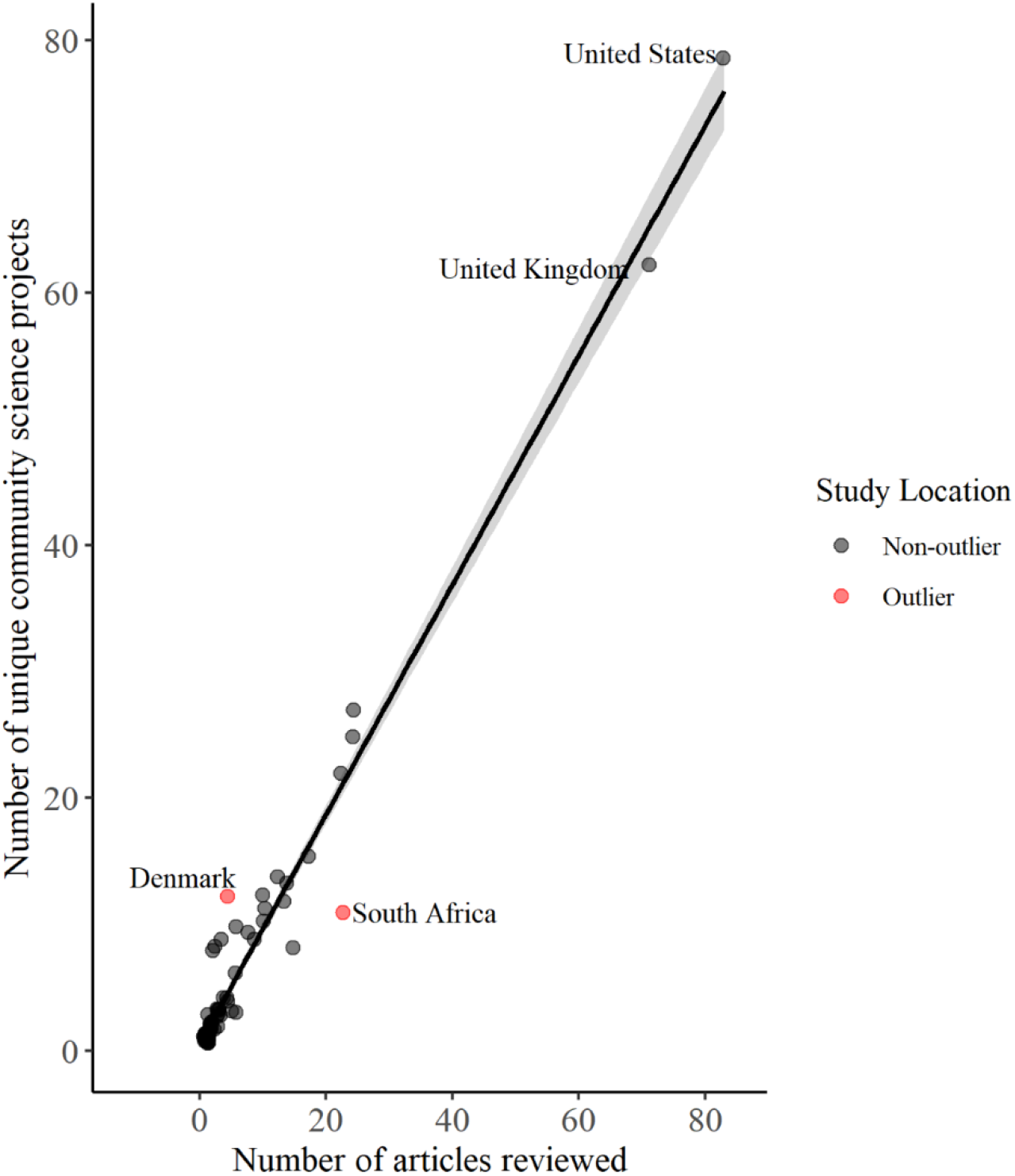
Linear relationship between the number of unique community science projects used in the scientific literature to monitor biodiversity in each country and the number of articles based in each country (β =0.91, R^2^ = 0.96, p = 3.36e-61). Data did not conform to normality assumptions, and were deliberately left untransformed, to examine patterns in residuals (see Appendix B; Fig. S3 for details). Note that we urge caution in interpreting between the bulk of points and the extreme values for the United States and the United Kingdom.

#### BOX 1. How community science is being used in the literature in Denmark and South Africa

**Figure 4.**
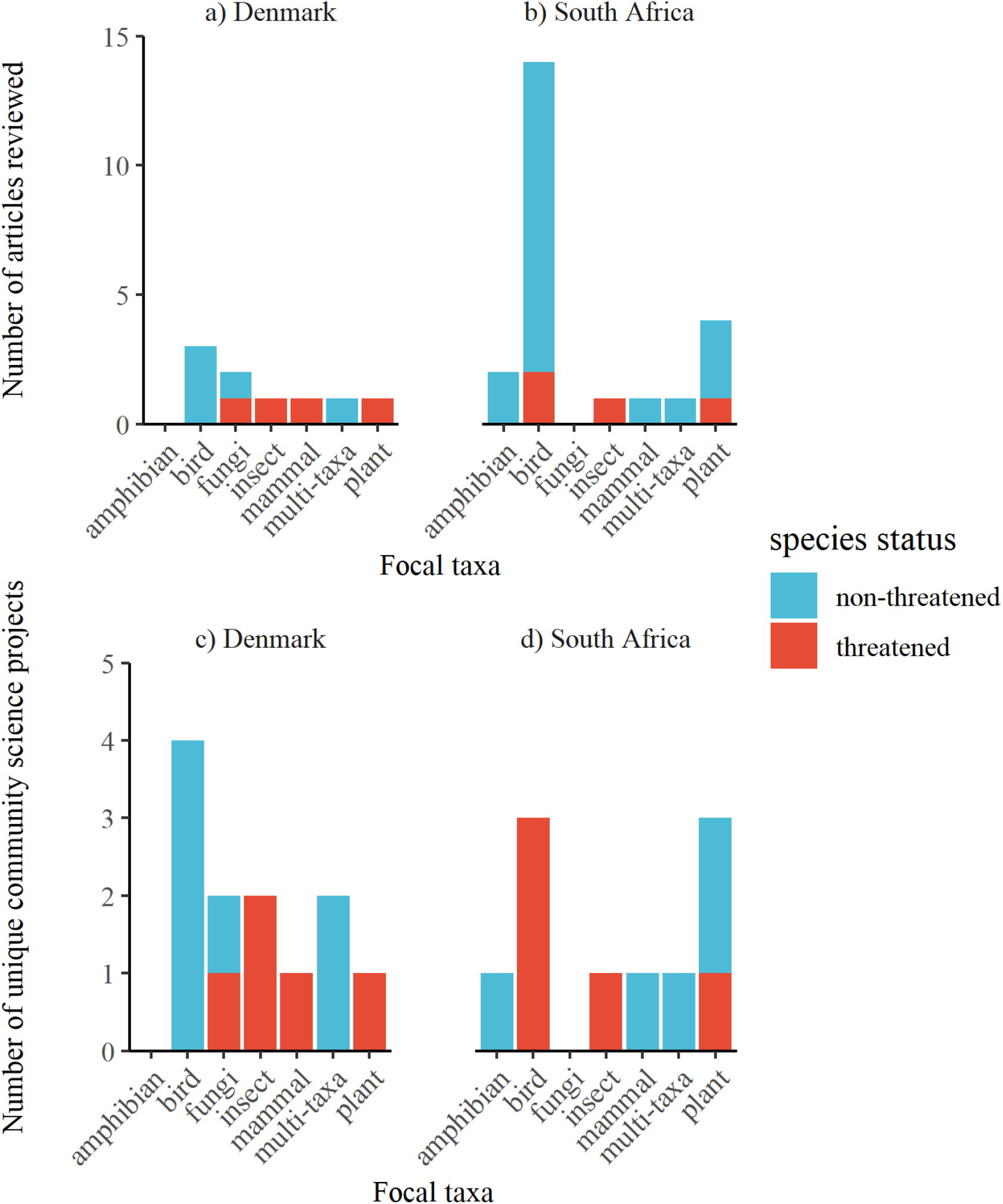
Taxonomic distribution within articles studying biodiversity in a) Denmark and b) South Africa. Taxonomic focus of the community science project for which data were used to study biodiversity in c) Denmark and d) South Africa. “Multi-taxa” denotes multiple taxonomic groups within a single article. A species status was noted as of conservation concern if it was characterized by the authors as such, inclusive of officially listed species and species demonstrating general decline that are not yet listed.

To gain insight into how the use of community science data in the literature can vary with the location of study, we examined the taxonomic distribution within the reviewed articles for two study locations with opposing tendencies: Denmark and South Africa (Fig. 3). Studies located in Denmark drew on a higher variety of community science projects than expected given the number of articles reviewed, while studies located in South Africa drew on a relatively low number of total community science projects given the high number of articles that studied South African biodiversity. Both studies located in Denmark and South Africa relied heavily on structured community science datasets: those in Denmark drew on a wide variety of atlases (general censuses of species’ distributions), and South African data stem largely from the South African Bird Atlas Project (SABAP), which was used for 14 of 23 total articles, and all but one of the studies on avian biodiversity.

In both countries, the types of projects favoured in our sample support the idea that baseline “high-quality” data from structured protocols, such as atlases and government-sponsored projects like SABAP and the BBS, are more readily incorporated into research than noisier datasets (Bayraktarov et al., 2019). This underscores the relevance of community science data for research and long-term monitoring in poorer countries (Callaghan et al., 2020), but also the usefulness of long-term monitoring projects for a greater variety of taxa in wealthier countries like Denmark. In any context, research can benefit from long-term community monitoring data only if the data are findable, accessible, and understandable (Theobald et al., 2015; Roche et al., 2021).

### 3.3. Conservation and environmental challenges addressed using community science in the literature

Sixty-three articles (approximately 18%) included at least one species that was considered of conservation concern, either described as “declining” by the authors or designated as “threatened”, “endangered” or a similar classification by a recognized conservation authority (Fig. 1). Most of these species were birds and invertebrates, reflecting the overall taxonomic pattern in our sample of the literature. The most common response metrics using community science data were distribution and range shifts, and abundance and/or trends (Fig. 5). A large proportion of articles (38%; n=127) from our literature search did not look at any specific threat to biodiversity. Articles that did examine a threat mainly focused on habitat loss, change or fragmentation (41.9%, n = 140).

**Figure 5.**
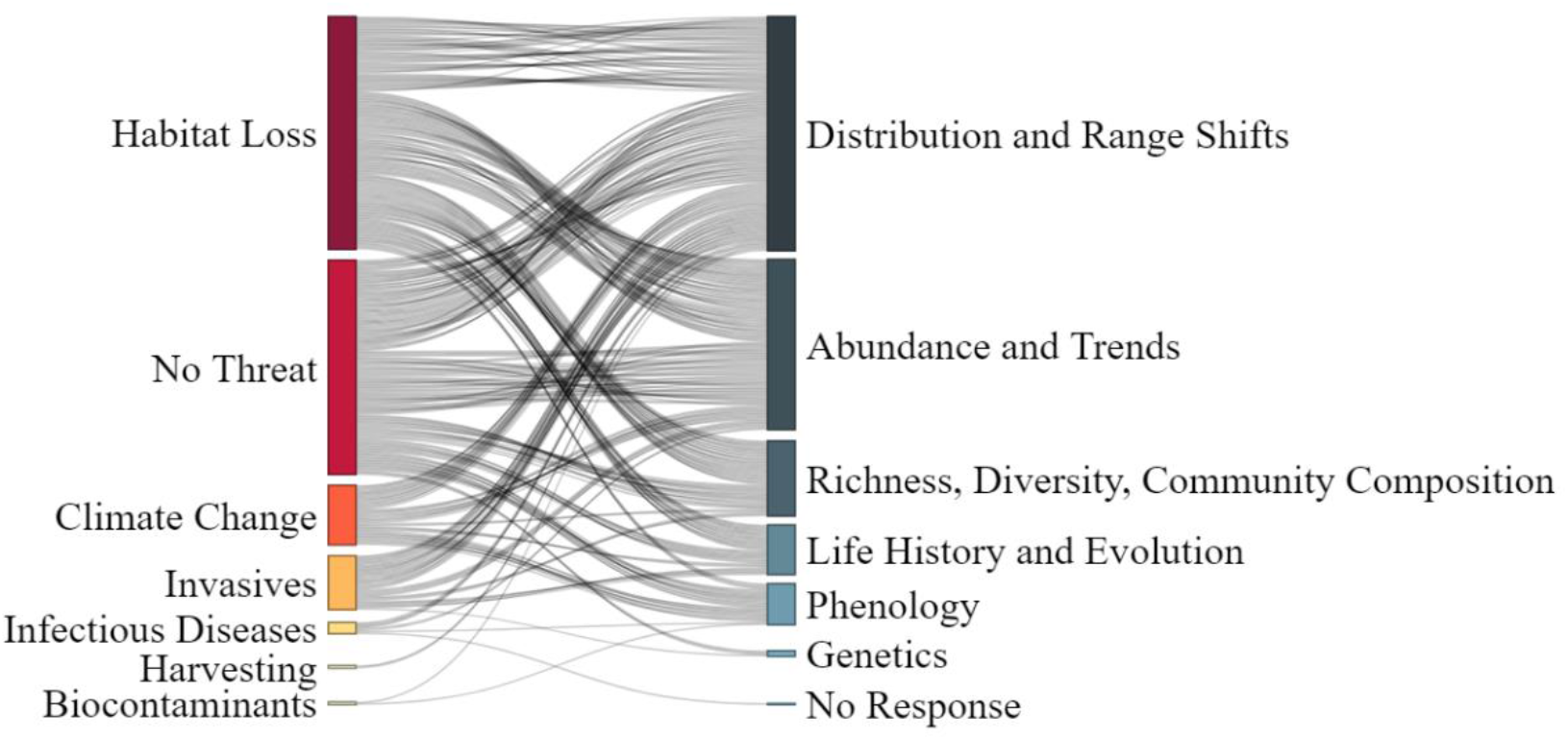
Type of threat examined, and the biological and ecological responses of biodiversity measured using community science data in the articles found in the literature search. Larger and darker rectangles represent a greater number of articles examining a given threat or response.

## 4. Discussion

Here we examined the patterns in community science data use in the published, peer-reviewed literature. We found that, in general, in a given location, more publications meant that a wider variety of projects were being used, but there were instances where certain countries were relying heavily on one well-vetted dataset. Additionally, the wealth of a country (as expressed by GDP per capita) explained very little of the variation in the use of community science data in publications. While the availability of data seemed to translate into more research on certain taxonomic groups such as birds and invertebrates, community science data are underutilized for other groups such as reptiles and amphibians, following the trends of conventional research biases (Donaldson et al., 2017). Finally, the use of community science to research at-risk species remains limited, and the links to specific threats are infrequently examined using these data.

These results support previous assertions that the uptake of community science data in the literature is driven in part by the range of the species and availability of data. Theobald et al. (2015) found that, although invertebrates are under-sampled compared to their overall abundance on earth, they are still the target of the greatest proportion of existing community science projects. Our results demonstrate that in this case, this greater data availability for invertebrates and birds translates into greater use in the published literature as well (Fig. 1). Although Theobald et al. (2015) found that birds had fewer community science projects dedicated to them than invertebrates, we found that the former were the subject of study more frequently than the latter. This discrepancy is understandable, given that birdwatching generates extensive data (Cordell & Herbert, 2002). Birds may also be more detectable compared to other taxonomic groups, given that they are often brightly coloured, highly vocal, and occupy a wide range of human-dominated habitats. These factors may play a role in the ability to monitor these species using community science efforts. Despite birds having fewer overall community science projects than invertebrates, there are disproportionately more projects collecting data on birds compared to the proportion of earth’s species that they represent (Theobald et al., 2015), and several of these are long-term monitoring programs that have existed for decades (e.g. The Breeding Bird Survey: Hudson et al. 2017) or even over a century (e.g. the Christmas Bird Count: Butcher 1990).

More general biases in conservation action, which in theory is informed by biodiversity monitoring research, may also explain why birds were the subject of study more frequently than invertebrates. Like birds, reptiles and amphibians also have a disproportionately greater number of community science projects dedicated to them compared to their relative abundance on earth (Theobald et al., 2015). However, when it comes to uptake in the literature, we find that publications making use of community science to monitor reptiles and amphibians are limited (Fig. 1), instead reflecting more conventional taxonomic biases in conservation research (Donaldson et al., 2017; Theobald et al., 2015). These biases are important to acknowledge and examine, as they are often propagated into conservation action and spending (Gordon et al., 2019).

Community science data are used for birds because they are widely available, but likely also because community science is often well-suited to monitoring such wide-ranging species. Community science is particularly used for large-scale monitoring (Chandler et al., 2017), and these large-scale projects are more likely to be used in published literature than smaller projects (Theobald et al., 2015). In addition to taxonomic biases, the relative lack of research on reptiles and amphibians may also be because such taxa tend not to be as wide ranging as birds or mammals, and therefore researchers do not need to rely on community science datasets in the same way they would for birds. This suggests that researchers turn to community science out of necessity; migratory birds can travel tens of thousands of kilometers throughout their annual cycle, making them challenging to monitor without relying on crowdsourced data.

Conservation research biases and conservation outreach usually disproportionally favour mammals (Donaldson et al., 2017), and research suggests that the proportion of mammal community science data reflects conventional researcher biases (Theobald et al., 2015). In our sample, mammals were less studied than birds and insects (Figure 1). This would imply that community science data are either not as well-suited for research on mammals as it is for birds, or that there is untapped potential for these data to be used in mammal conservation research.

The cause of the geographic patterns of community science use found in our review are difficult to discern; they could be merely a reflection of publication rates varying among countries, or English language bias, but also the availability of community science data, as suggested by Theobald et al. (2015), which is beyond the scope of this study. Nonetheless, these geographic patterns are interesting for several reasons. It could be that developing countries rely more on community efforts to monitor biodiversity given sparse funding for professional monitoring. In contrast, while more cost-efficient at larger scales (Heigl et al., 2017), community science programs are certainly not free, and issues with governance and safety could limit the persistence of these projects (Pocock et al., 2019). Additionally, community scientists in wealthier countries may have more free time at their disposal to contribute to community science projects, thus generating more data available to be used in research and consequently more publications.

For endangered species, community science projects must contend with special considerations about project design, volunteer engagement, and open-data practices, where the safety of imperiled species is of particular concern (Lindenmayer and Scheele 2017; Lennox et al. 2020). Common species are also inherently easier to monitor and study than those that are rare. It is therefore unsurprising that the bulk of the literature in our sample examines species that are not at risk (Fig. 1). The exception here was with fungi, almost all of which were described as at risk or declining. Given the difficulty detecting and identifying species of fungi, and the relative lack of interest in them compared to other taxa, it is possible only the most interesting and/or threatened species are reported and researched.

Our review also highlighted that community science efforts were largely used to measure abundance, trends, distributions, and range shifts. This finding was expected given the ability of community science to provide the extensive data necessary for studying these topics (Dickinson et al., 2010; Lin et al., 2022; Theobald et al., 2015). However, for similar reasons, community science data can be well suited to measuring changes in phenology (Binley et al., 2021), yet we found limited evidence of this application in our sample of the literature. Other responses such as collecting genetic samples may simply be limited because this is a novel and developing field (Granroth-Wilding et al., 2017).

Surprisingly, relatively few studies used community science data to examine the effects of climate change and invasive species on biodiversity. Given that these are threats that often occur at broad spatial scales, it would be advantageous to use community science to measure their impacts (Binley et al., 2021). However, community science may not be the best option for measuring threats of direct exploitation. Community science is inherently open and sharing the location of these species may put them at greater risk (Tulloch et al., 2018). Safeguards that protect the location of at-risk species and the well-being of wildlife populations, as well as the safety of the people observing these species, can be put in place for monitoring sensitive species with community science initiatives (Soroye et al., 2022).

We also recognize a gap whereby community science could be used more often to examine threats. Although most community science programs are often better suited for baseline surveillance monitoring (Dickinson et al., 2010), these data can still be used to test hypotheses (Yoccoz et al., 2001). Community science programs are frequently designed with a particular goal in mind, such as measuring population trends (e.g., the Breeding Bird Survey; Hudson et al. 2017), changes in phenology (e.g., Plant Watch; Gonsamo, Chen, and Wu 2013), or monitoring a specific threat (e.g., Fitzgerald et al., 2014). Often, the proponents of such programs are under the impression that these data are then used in scientific research, which unfortunately is not always the case (Theobald et al. 2015). Given the urgency of the biodiversity crisis, we expected that community science projects would be used more in the literature to investigate specific threats and their impacts on biodiversity (Buxton et al., 2021). Thus, it appears that conservation research is not making full use of community science data to link known threats with biological and ecological responses, despite the evident potential to do so. This unfortunately may reflect a general pattern in the literature, whereby research effort often neglects actionable and high-priority questions (Buxton et al., 2021). Although we acknowledge that our evidence synthesis examines only a representative sample of the literature, rather than systematically reviewing it, we believe there is room for more applied research using community science to address threats to biodiversity, particularly those operating at larger scales. Indeed, one of the strengths of community science is that conservation practitioners and researchers often have these data readily available, circumventing the time and expense needed to collect further information needed to answer pressing questions when it is not always warranted to do so.

Community science is a growing source of global biodiversity data that is particularly useful for covering large spatial and temporal extents, and is generally more easily available and accessible than other data sources, such as grey literature or “file-drawer” data (Haddaway & Bayliss, 2015). Our results suggest that there is untapped potential for community science data use, in examining broader sets of taxa, and in tracking the effects of biodiversity threats such as climate change and invasive species. Our results also suggest that key lessons can be learned from certain jurisdictions that make extensive use of community science data for biodiversity monitoring. While inclusion in peer-reviewed publications may not be the ultimate goal of community science projects (nor necessarily should it be), community science project managers can often be under the impression that their work is being used in the literature, even if this is not necessarily the case (Theobald et al., 2015), and there are a multitude of applications that are often underutilized (Binley et al., 2021). Future interdisciplinary research could examine the social, political, and economic conditions necessary for successfully implementing a wide variety of community science programs in different countries, and the associated likelihood of community science data uptake within peer-reviewed literature and beyond.

## 5. Glossary

Community science data: Data collected through public participation in surveys and monitoring. Participants can be experts or novices, and data collection protocols can be unstructured (fully opportunistic), semi-structured (largely opportunistic but including the collection of covariate data) or structured (following a defined protocol). Participants are volunteers and are not paid.

## 6. Acknowledgements

We would like to thank the many talented and dedicated community scientists that make our research possible.

## Notes

https://osf.io/wze3a/?view_only=524f5be46c0b467ca16061482b0c8465

### Competing Interest Statement

The authors have declared no competing interest.

