## Supplemental materials for "Patterns of community science data use in peer-reviewed research on biodiversity"

Supplementary Materials:

APPENDIX A: Literature Search and Data Extraction

WoSCC Institution subscriptions:

- Science Citation Index Expanded (1900 - present)
- Social Sciences Citation Index (1956 - present)
- Arts & Humanities Citation Index (1975 - present)
- Conference Proceedings Citation Index - Science (1990 - present)
- Conference Proceedings Citation Index - Social Science and Humanities (1990 - present)
- Book Citation Index - Science & Social Science (2008 - present)
- Current Chemical Reactions (2008 - present)
- Index Chemicus (2008 - present)
- Part of the larger [Web of Science](https://library.carleton.ca/find/databases/web-knowledge)

Scopus Institution subscriptions: Full database

Data coding and extraction

For each study, we grouped the threat (explanatory) and response variables into categories, based on Dickinson et al., 2010) (Table S1). We also noted when studies did not examine a particular threat (e.g., building species distribution models) nor a species response metric (e.g., studying disease spread).

Table S1. Description of the explanatory (threat) and response variables examined in each study from the literature search, based on Dickinson et al., (2010).

| **Variable** | **Metric** | **Description** |
| --- | --- | --- |
| **Explanatory** | Climate change | Examines threats related to climate variables, or changes in climate over time |
|  | Habitat loss, change or fragmentation | Examines habitat loss, fragmentation of habitat, or conversion of habitat to another land cover type |
|  | Infectious disease | Tracks infectious disease spread in one or more species |
|  | Biocontaminants | Assesses environmental contaminants such as heavy metals or acid deposition |
|  | Invasive species | Investigates species occurring outside of their native range that have a negative impact on native species |
|  | Harvesting | Species are harvested for human use or consumption, through legal or illegal means |
| **Response** | Distribution and range shifts | Assesses species distributions or ranges, including changes or trends |
|  | Phenology | Measures any temporal aspect of a species life cycle |
|  | Richness, diversity, community composition | Any self-described measure of species richness, diversity or community composition, including turnover |
|  | Abundance and trends | Estimates of species abundance, or change in abundance over time |
|  | Life history and evolution | Assesses ecological strategies for survival or reproduction, and changes in these strategies over time |
|  | Genetics | Collects genetic information on species, either to specifically assess genetic makeup and variability or as a means of measuring any of the other response variables (e.g. distribution) |

APPENDIX B: Further methods and Supplementary Figures

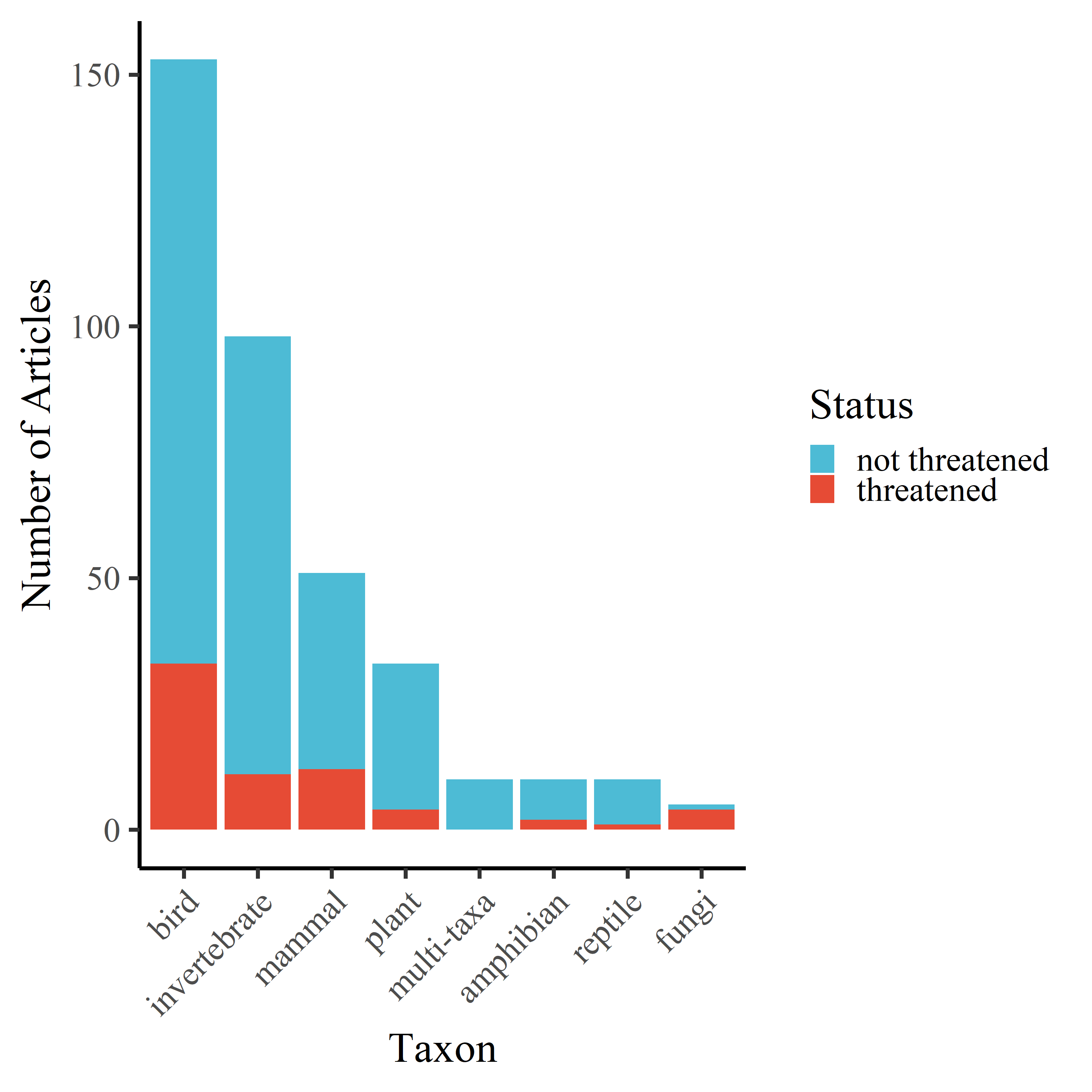

Figure S1. The number of articles in the selection of articles captured by the literature review that studied each taxonomic group. Blue represents the proportion of articles that studied species that are described as being “threatened” by the author of the article, either due to population declines or because they are officially listed under some conservation authority as being at risk or in decline. “Multi-taxa” denotes an article or project that includes multiple taxonomic groups within a single study, and “threatened” versus “not threatened” in this case refers to whether any of the species examined were described as at risk.

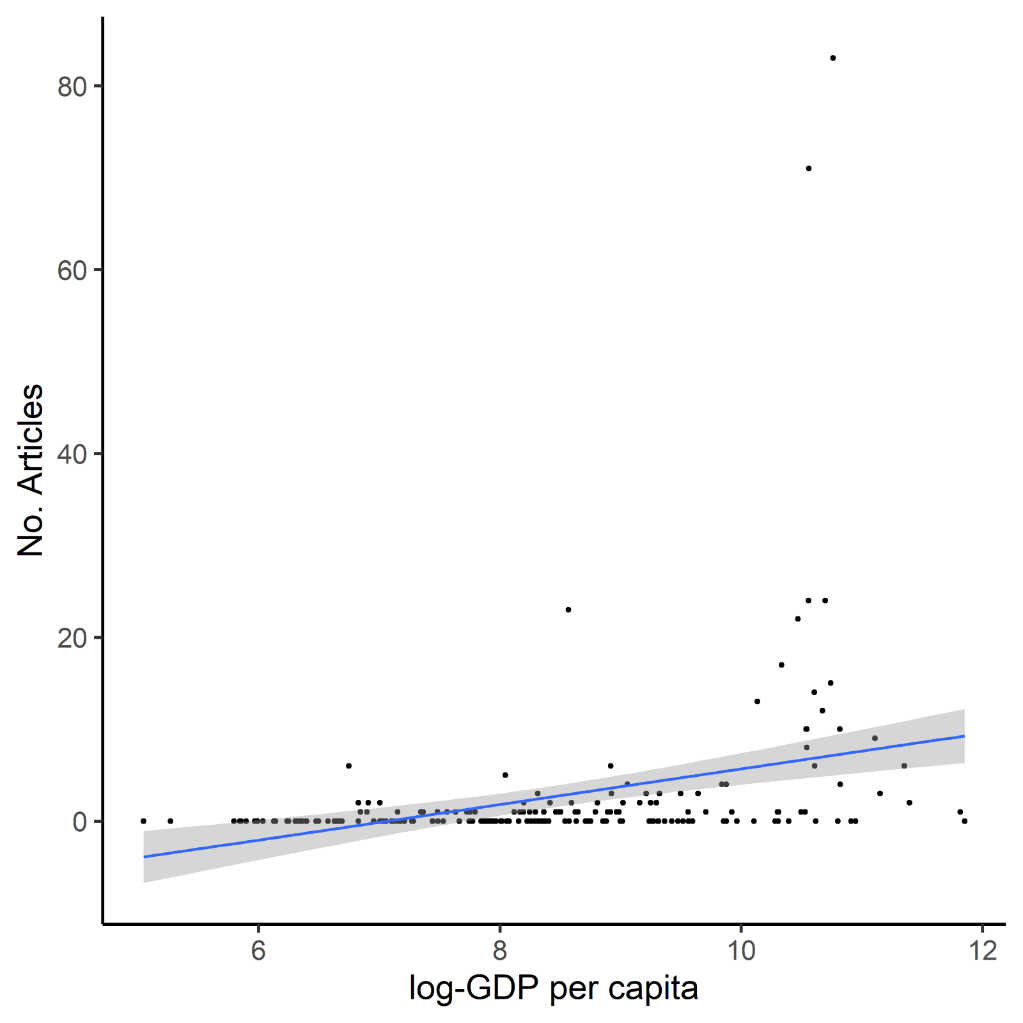

Figure S2. Log-linear relationship between GDP per capita and the number of articles in each country making use of community science data for biodiversity (β = 1.936, p < 0.0001, R^2^ = 0.108).

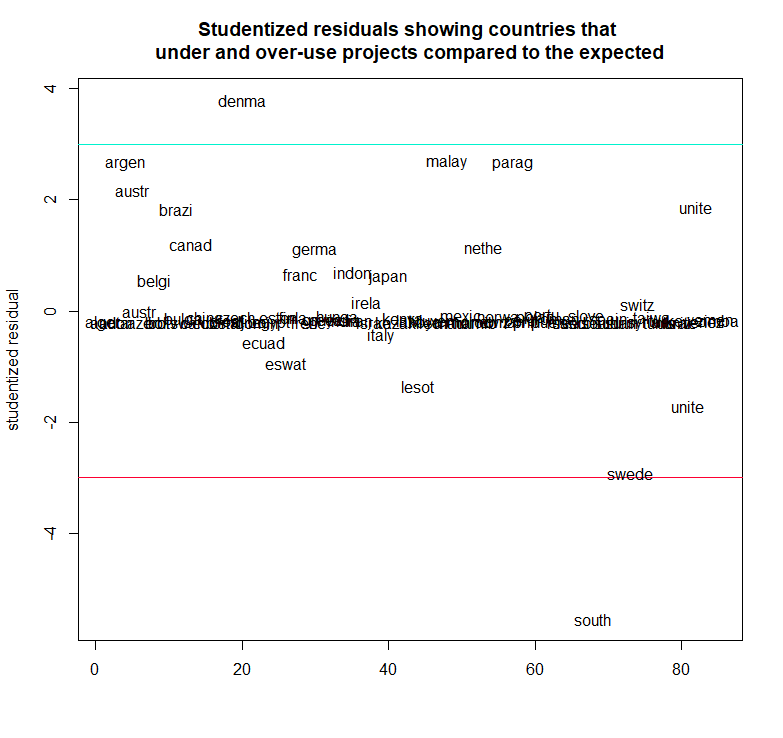

Figure S3. Studentized residuals showing countries that under- and over-used projects compared to the expected linear relationship. Two countries diverged from the expected linear relationship between the number of articles and the number of distinct projects: for the number of articles included in our review, we found fewer distinct projects than expected in South Africa, and more projects than expected in Denmark.
